# Modeling Stimulus-Induced Stress Responses in Microglia-like Cells Using a Commercial iPSC-dCas9-KRAB Line

**DOI:** 10.1101/2025.08.30.673269

**Authors:** Devin M. Saunders, Nicola A. Kearns, Bernard Ng, Faraz Sultan, Himanshu Vyas, Ricardo A. Vialle, Eric M. Clark, Sashini De Tissera, Jishu Xu, David A. Bennett, Yanling Wang

## Abstract

Commercially available human iPSC lines with inducible CRISPR interference (CRISPRi) systems offer scalable platforms for gene function studies. One widely available line, the AICS-0090 dCas9-KRAB iPSC line developed by the Allen Institute, has been extensively validated for genomic integrity and stem cell potency. However, its utility in modeling specialized immune cell types such as microglia — and in assessing their functional responses to disease-relevant stimuli — has not been fully established. Here, we evaluated the AICS-0090 line for its ability to differentiate into microglia-like cells, support efficient gene knockdown, and respond to environmental stressors. We assessed its differentiation capacity by qPCR, flow cytometry, and immunocytochemistry, confirming reproducible expression of microglial surface markers at early and late timepoints. Gene knockdown efficiency was validated both at the single-gene level and in pooled CRISPRi screens. Focusing on functional responsiveness, we exposed the microglia-like cells to two distinct stimuli: amyloid-β (Aβ), a disease-associated trigger in Alzheimer’s disease, and lipopolysaccharide (LPS), a classical inflammatory signal. Transcriptomic and functional analyses revealed stimulus-specific responses: Aβ induced limited activation of stress and inflammation pathways, whereas LPS elicited broader transcriptional reprogramming and cytokine release. Signatures from both conditions partially overlapped with ex vivo human and mouse microglial states. Together, these findings support the use of the AICS-0090 dCas9-KRAB iPSC-derived microglia-like cells as a flexible and tractable model for gene function interrogation under defined inflammatory contexts, with potential for future applications in neuroimmune modeling and perturbation-based screening.

## Introduction

Commercially available human iPSC lines with inducible CRISPR interference (CRISPRi) systems offer scalable platforms for gene function studies. One such line, the AICS-0090 dCas9-KRAB iPSC line developed by the Allen Institute, has been validated for genomic integrity, pluripotency, and differentiation into multiple lineages, as documented in the publicly available [Certificate of Analysis (CoA)](https://www.coriell.org/0/PDF/Allen/ipsc/AICS-0090-391_CofA.pdf). However, its utility in modeling specialized cell types such as microglia — and in assessing their transcriptional and functional responses to environmental challenges — has not been fully validated.

Microglia are the brain’s resident immune cells and key sensors of the central nervous system microenvironment (Frank et al., 2019; Colonna and Butovsky, 2017). They maintain homeostasis and rapidly respond to diverse perturbations. When dysregulated, microglial activity contributes to neuroinflammation and neurodegenerative processes, including Alzheimer’s disease (AD)(Deczkowska et al., 2018; Gao et al., 2023; Hickman et al., 2018). A better understanding of how human microglia respond to external stimuli is essential for dissecting disease mechanisms and modeling injury- or aging-associated states in vitro.

Stress responses are core cellular programs that enable cells to cope with physiological and pathological challenges. Among AD-related stressors, amyloid-β (Aβ) peptides act as chronic signals that induce innate immune activation (Deczkowska et al., 2018; Heneka et al., 2015). In contrast, lipopolysaccharide (LPS), a potent exogenous stimulus, elicits robust inflammatory responses(Hanisch and Kettenmann, 2007; Kettenmann et al., 2011). While both are widely used to model microglial activation, the extent to which they trigger overlapping versus distinct molecular and phenotypic programs remains incompletely defined.

To address this, we utilized the AICS-0090 dCas9-KRAB iPSC line to model stimulus-induced stress responses in microglia-like cells (iMGLs). We confirmed its ability to differentiate into iMGLs, support effective gene knockdown using single-gene and pooled CRISPRi screens, and respond transcriptionally and functionally to Aβ and LPS stimulation. This work supports the AICS-0090 iMGL platform as a flexible and tractable system for modeling microglial responses to environmental stimuli and genetic perturbations, laying the foundation for future studies exploring gene-environment interactions in neuroimmune contexts.

## Results

### Characterizing the dCas9-KRAB iPSC line in microglia-like cell differentiation and gene perturbation

To evaluate the utility of the dCas9-KRAB human iPSC line for generating iMGLs, we first assessed its differentiation capacity using published protocols (Abud et al., 2017; McQuade et al., 2018) (Fig. 1A). Brightfield imaging showed morphological changes during the differentiation timeline, with day 40 iMGLs appearing more branched compared to earlier time points (Fig. 1B). We evaluated gene expression of microglia-associated genes *APOE*, *CD33*, and *TREM2* by qRT-PCR at key time points (iPSCs, day 12, day 24, and day 40). These genes showed variable expression patterns over time, with *CD33* and *TREM2* detectable at day 12 and increasing by day 24, while *APOE* expression peaked at day 24 before declining slightly by day 40 (Fig. 1C). Flow cytometry analysis demonstrated robust CD43 expression at day 12, indicative of hematopoietic progenitor cell (iHPC) generation, and early CD45 expression. By day 40, iMGLs exhibited strong CD45 positivity and expressed canonical microglial surface markers including CD11b, CX3CR1, TREM2, P2RY12, PU.1, IBA1, and TMEM119 (Fig. 1D). Immunofluorescence imaging confirmed the expression of several of these markers at the protein level (Fig. 1E). Finally, functional phagocytosis assays using pHrodo-labeled Aβ demonstrated that approximately 40% of iMGLs exhibited Aβ uptake activity (Fig. 1F).

**Figure 1.**
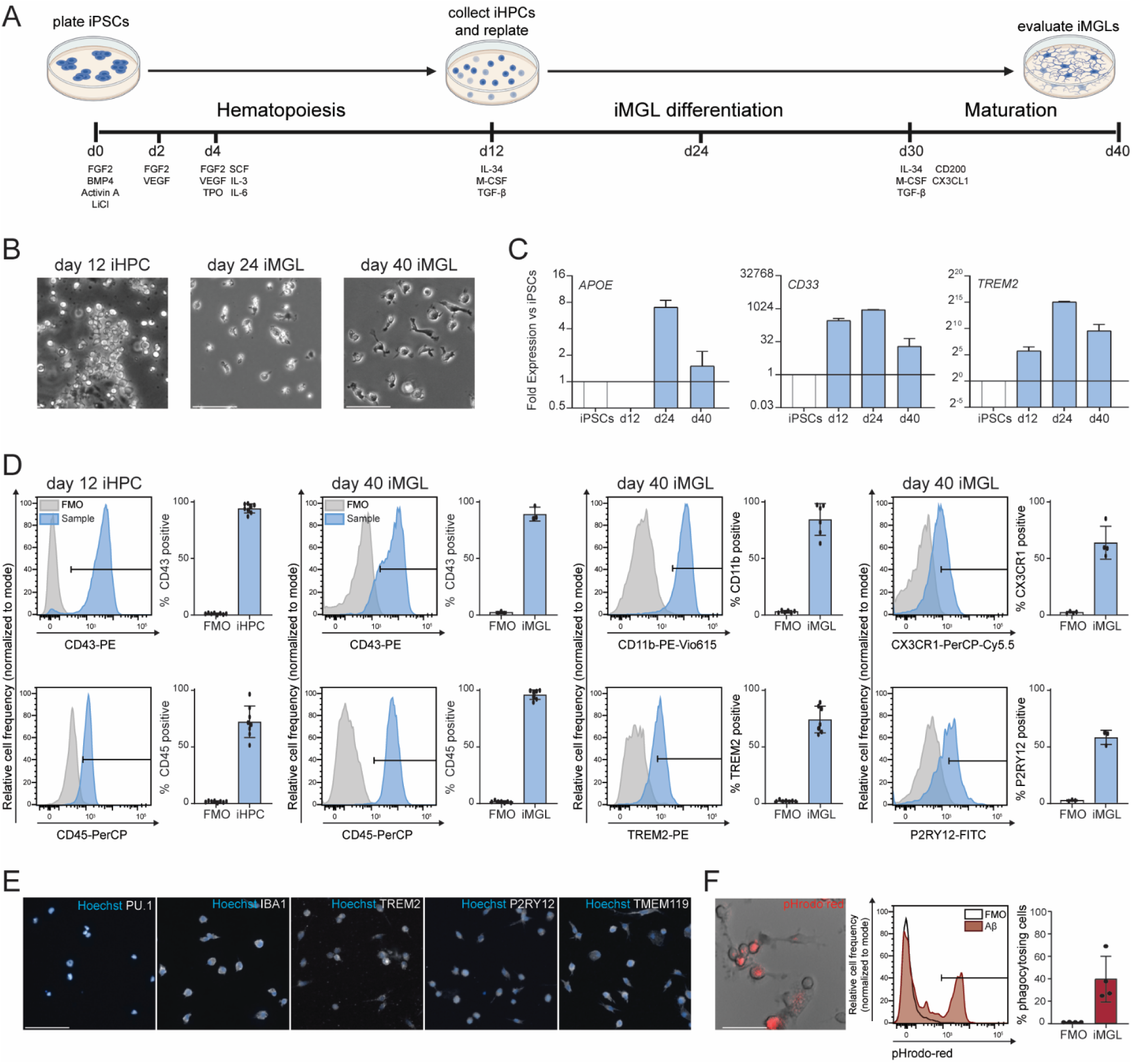
Differentiation and characterization of dCas9-KRAB iMGL. (A) Schematic illustrating a modified iPSC-derived iMGL differentiation protocol. dCas9-KRAB iPSCs are differentiated to iHPCs for 12 days and then to iMGL for an additional 18 days, after which they’re matured for 10 days. (B) Representative brightfield images of iHPCs at differentiation day 12 and iMGLs at differentiation days 24 and 40. Scale bar: 50 µm. (C) Quantitative RT-PCR analysis of APOE, CD33, and TREM2 expression across multiple differentiation stages, demonstrating transcriptional maturation of iMGLs. (D) Flow cytometry analysis of CD43 and CD45 expression on differentiation days 12 and 40, plus additional maturation markers on day 40 including CD11b, CX3CR1, TREM2, and P2RY12. Representative histograms and quantification illustrate robust expression of myeloid markers during iMGL differentiation. Data represent 3-8 independent iMGL differentiations (mean ± SD). (E) Immunocytochemistry (ICC) images show iMGL express canonical microglial markers at day 40, including PU.1, IBA1, TREM2, P2RY12, and TMEM119. Scale bar: 50 µm. (F) Phagocytosis assay in live dCas9-KRAB iPSC-derived iMGLs stimulated with Aβ. Representative brightfield and fluorescent image of actively phagocytosing iMGLs with internalized pH-sensitive pHrodo Red-labeled Aβ, which fluoresces red within acidic phagosomes. Histogram and quantification illustrate pHrodo Red-positive cells relative to FMO controls. Scale bar: 50 µm. Data represent 4 independent iMGL differentiations (mean ± SD).

To evaluate the gene perturbation capacity of the dCas9-KRAB iPSC line, we first targeted CD33, an Alzheimer’s disease risk gene highly expressed in microglia (Fig. 2A). Lentiviral vectors expressing sgRNAs and GFP reporters were transduced into day 30 iMGLs. By qRT-PCR, we observed partial CD33 knockdown, with guideRNA2 producing the strongest reduction in transcript levels (Fig. 2B). This knockdown was further confirmed at the protein level by flow cytometry, which showed decreased CD33 surface intensity specifically in cells transduced with guideRNA2 (Fig. 2C), validating the effectiveness of the CRISPRi machinery in this system. To assess scalability, we performed a pilot pooled CRISPRi screen targeting 165 AD-related genes (sFig.1). sgRNAs were cloned into lentiviral vectors and transduced into iPSCs prior to differentiation (Fig. 2D). sgRNA representation was quantified across the differentiation timeline by next-generation sequencing at days 0, 12, 24, 36, and 45. As shown in Fig. 2E, the initial sgRNA distribution followed a bell-shaped histogram, consistent with even representation at the iPSC stage. We then compared dropout rates between sequential timepoints (d24 vs. d12, d36 vs. d24, d45 vs. d36) to identify survival-essential genes. This analysis revealed multiple genes—including *TGFBR2, TOMM40L*, *ABCA1* and others—whose depletion significantly impacted cell survival (Fig. 2F), aligning with known regulators of microglial biology and AD pathogenesis (Butovsky et al., 2013; Chen et al., 2021, 2023; Karasinska et al., 2013; Zöller et al., 2018).

**Figure 2.**
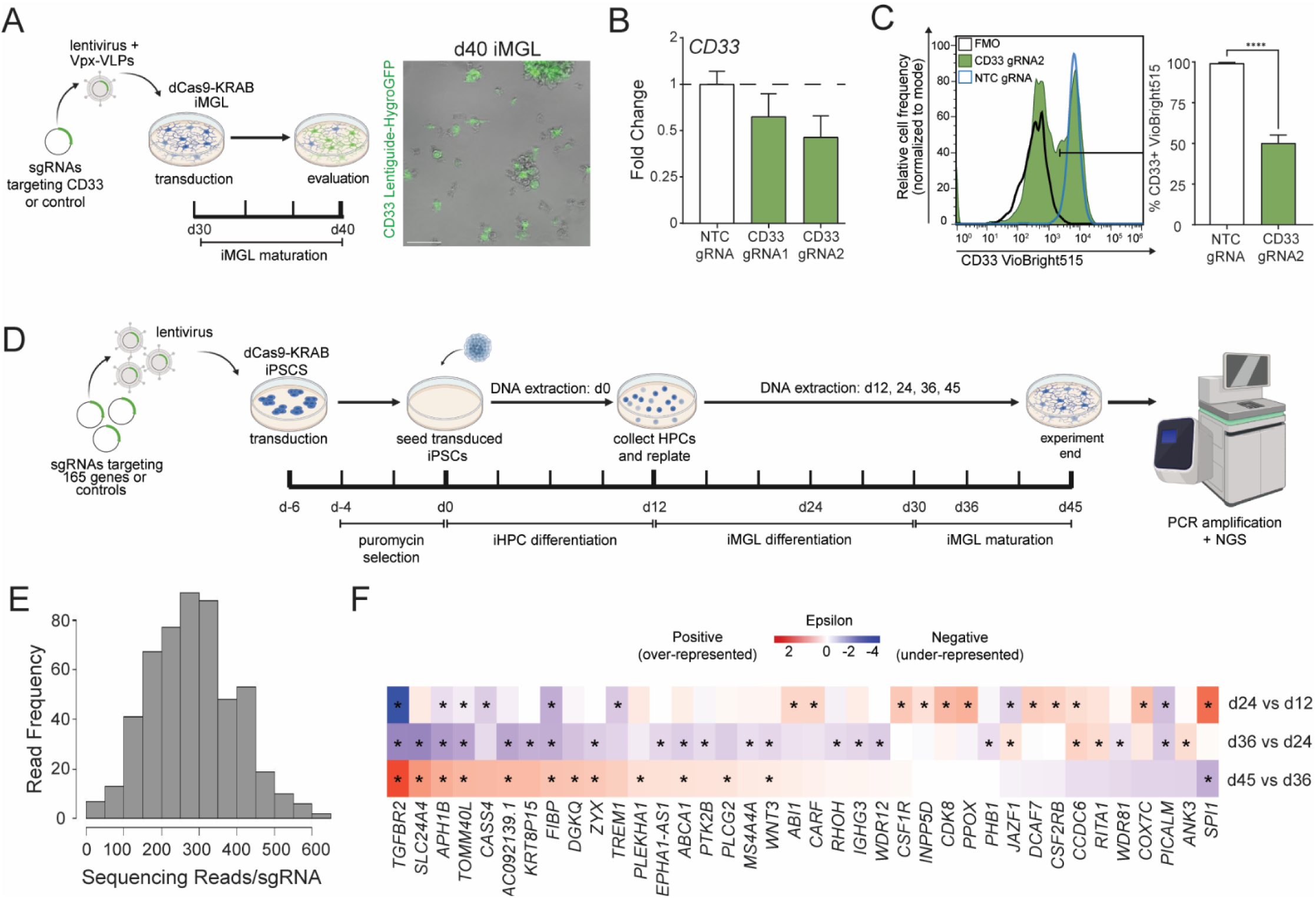
dCas9-KRAB iMGLs retain CRISPRi functionality after differentiation for both individual gene knockdown and pooled screens. (A) Schematic of viral delivery of CD33 or non-targeting control (NTC) eGFP-tagged gRNAs for CRISPRi knockdown in dCas9-KRAB iMGL using lentivirus combined with viral-like particles (VLPs) carrying Vpx protein (Vpx-VLPs). Representative brightfield and fluorescence image of day 40 iMGLs shows successful delivery of the CD33 gRNA, indicated by green fluorescence. Scale bar: 50 µm. (B) Quantitative RT-PCR analysis of CD33 expression following transduction with NTC or two independent CD33 gRNAs. Data represent fold change knockdown of CD33 relative to NTC. (C) Flow cytometry analysis of CD33 protein levels in day 40 iMGLs, 10 days after transduction with NTC or gRNA2. Representative histogram and quantification demonstrate efficient knockdown of CD33 protein in gRNA2-transduced cells. Data represent fold change knockdown of CD33 relative to NTC across 4 independent iMGLs differentiation (mean ± SD). (D-E) Schematic of pooled CRISPRi screen workflow in dCas9-KRAB iMGLs. Cells were transduced with a 165-gene sgRNA library or controls, selected with puromycin, differentiated to iMGL, and collected for DNA extraction across multiple differentiation stages. PCR amplification and NGS were used to quantify sgRNA abundance and distribution. (F) Heatmap of significant gene-level enrichment (Epsilon), showing over- or under-representation of sgRNAs between successive differentiation stages. *\*p* < 0.05.

### Aβ and LPS elicit robust morphological and transcriptional responses in dCas9-KRAB iMGLs

To validate the utility of the dCas9-KRAB iMGLs for modeling microglial responses to pathological stimuli, we stimulated cells with either oligomeric Aβ (2 μg/mL) or LPS (1 μg/mL) and assessed morphological and transcriptional changes. Morphologically, both stimuli triggered activation signatures with increased cellular processes and enlarged somas compared to unstimulated cells, as observed by immunofluorescence imaging for IBA1 and CD45 (Fig. 3A). Notably, LPS treatment induced a more hypertrophic, multipolar morphology and a pronounced redistribution of CD45 to the plasma membrane, reflecting a classical pro-inflammatory activation state (Fig. 3A).

**Figure 3.**
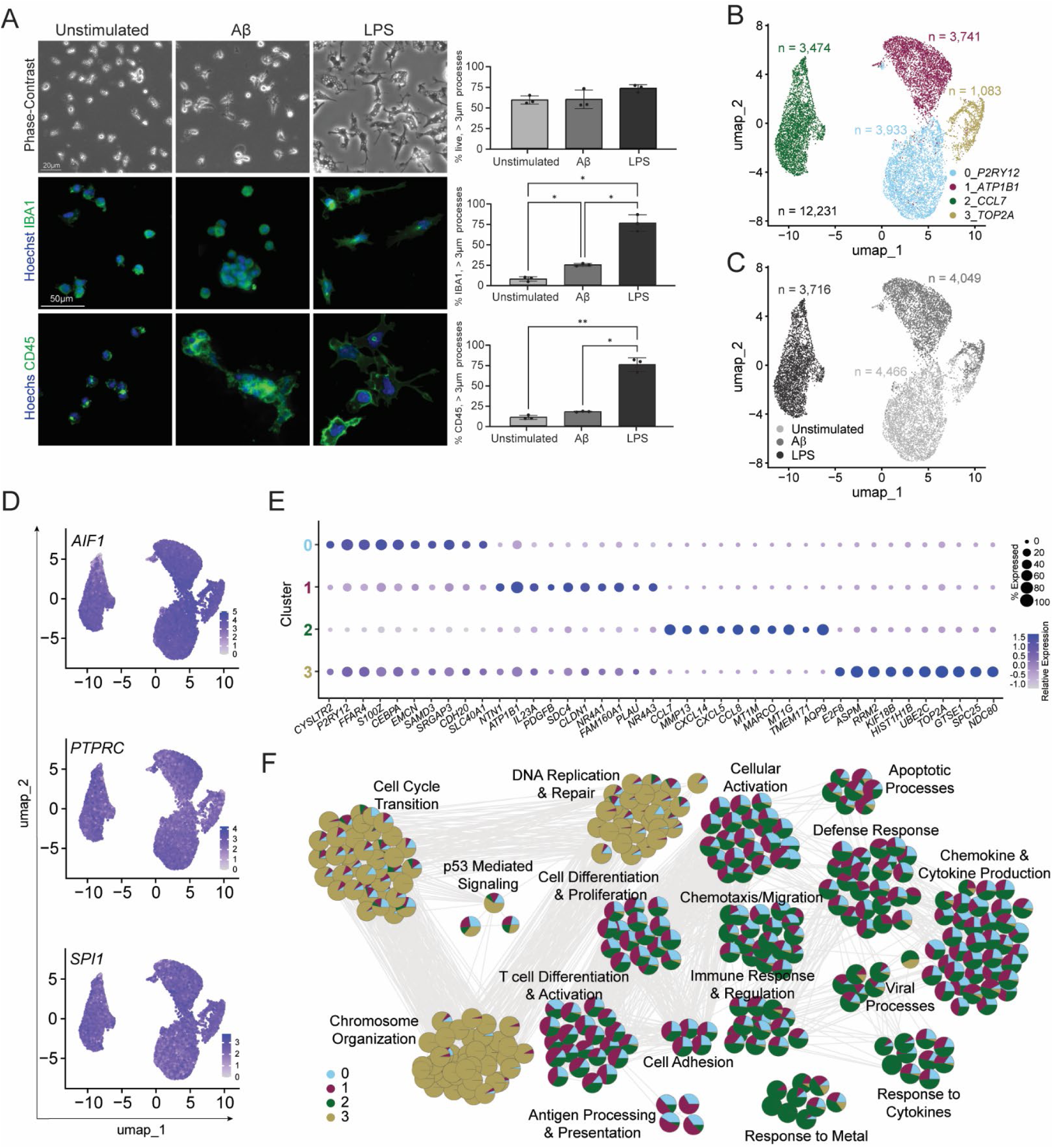
Aβ and LPS elicit distinct morphological phenotypes and transcriptional states in iMGLs. (A) Representative phase-contrast and IBA1/CD45 immunofluorescence images showing distinct morphological phenotypes in unstimulated, Aβ-treated, and LPS-treated iMGLs. Scale bars: brightfield 20 µm, IBA1/CD45 50 µm. Data represent 3 independent iMGL differentiations, with ∼900–2,000 cells quantified per condition across imaging modalities (mean ± SD). *\*p* < 0.05; \*\**p* < 0.01 (repeated-measures one-way ANOVA with Tukey’s multiple-comparisons test). (B-C) UMAP visualization of 12,231 high-quality iMGLs profiled by scRNA-seq, revealing segregation into four major transcriptional clusters largely segregated by condition. (D) Feature plots showing expression of canonical microglial identity genes (*AIF1*, *PTPRC*, and *SPI1*) across all clusters, confirming preservation of microglial lineage identity following stimulation. (E) Dot plot displaying the top 10 differentially expressed genes for each transcriptional cluster, ranked by average log_2_ fold change. (F) Gene Ontology (GO) enrichment analysis performed on the top 200 differentially expressed genes per cluster, highlighting enriched biological process, molecular function, and cellular component terms associated with cluster-specific transcriptional programs. Complete gene lists and enrichment results are provided in Table S3.

We next evaluated the transcriptional responses of the dCas9-KRAB iMGLs using single-cell RNA sequencing (scRNA-seq) across control, Aβ-treated, and LPS-treated cultures. Following data generation, we applied quality control filtering based on metrics such as total gene count, UMI count, mitochondrial transcript percentage, and ribosomal content (sFig. 2A). After dimensionality reduction and clustering, four major transcriptional clusters emerged that were primarily segregated by treatment condition (Fig. 3B–C). All clusters retained expression of core microglial markers such as *AIF1*, *PTPRC*, and *SPI1(*Jurga et al., 2020), supporting microglial identity (Fig. 3D). Examination of the top differentially expressed genes (DEGs) revealed that each cluster exhibited a distinct transcriptional signature (Fig. 3E; sFig. 2B–C), consistent with treatment-specific responses. Cluster 0 contained predominantly unstimulated iMGLs and was enriched for canonical homeostatic markers such as *P2RY12, FCRL4*, and *CEBPA* (Gómez Morillas et al., 2021; van Wageningen et al., 2019; Li et al., 2023; Wang et al., 2024). Cluster 1 was enriched in Aβ-treated cells and showed elevated expression of genes related to lipid metabolism and immune activation, including *SC4D, NR4A1/3,* IL23A (García-Domínguez, 2025; Phelan et al., 2021; Rothe et al., 2017). Cluster 2, dominated by LPS-treated cells, expressed genes involved in classical inflammatory and interferon-associated responses, including *CCL7, MMP13*, and *MARCO* (Areschoug and Gordon, 2009; Könnecke and Bechmann, 2013; Xue et al., 2021). Cluster 3 comprised proliferative cells found in the unstimulated and Aβ conditions and was marked by cell cycle–related genes such as *TOP2A, E2F2*, and *UBE2C* (Filipovich et al., 2025).

Gene ontology analysis of the top 200 genes per cluster revealed functional programs linked to each condition (Table S3). Aβ exposure was associated with upregulation of genes involved in antigen processing and T cell differentiation, whereas LPS-treated cells showed strong enrichment for chemotaxis, cytokine signaling, and defense response pathways. A subset of cells from both stimuli conditions displayed a proliferation signature with elevated expression of cell cycle regulators (Fig. 3F).

Together, these results demonstrate that dCas9-KRAB iMGLs mount distinct morphological and transcriptional responses to Aβ and LPS, capturing stimulus-specific activation states.

### Differential gene expression analysis reveals shared and divergent transcriptional responses in Aβ- and LPS-stimulated dCas9-KRAB iMGLs

To systematically compare the transcriptional impact of Aβ and LPS stimulation in dCas9-KRAB iMGLs, we first performed differential gene expression analysis against untreated controls. LPS stimulation elicited broader and more robust transcriptional shifts compared to Aβ exposure, with a greater number and magnitude of differentially expressed genes (Fig. 4A-B; Table S4). These results suggest that LPS triggers a more extensive reprogramming of the iMGL transcriptome, consistent with a stronger inflammatory activation state.

**Figure 4.**
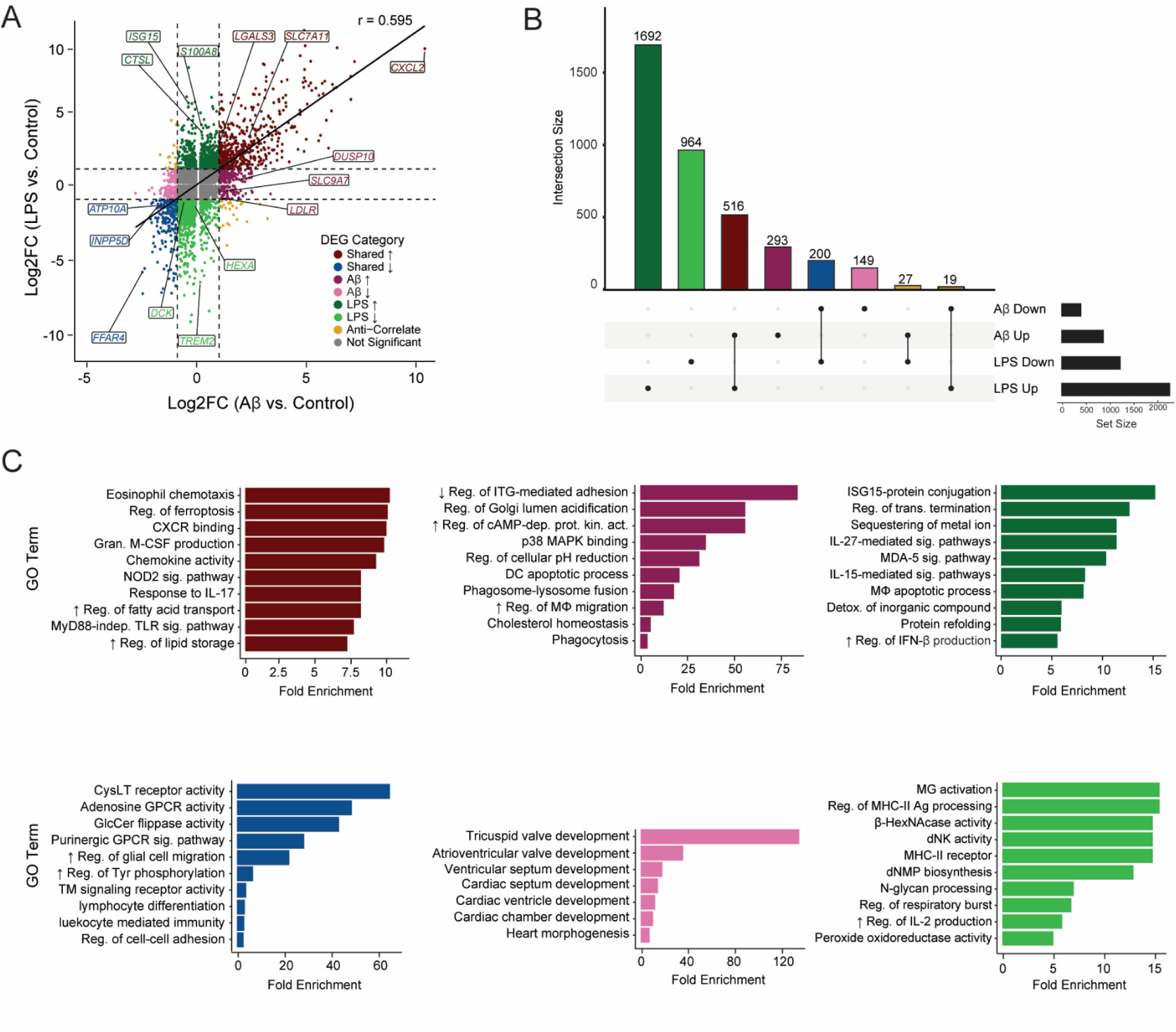
Aβ and LPS induce shared and distinct transcriptional stress responses. (A) Differential gene expression profiles in Aβ- and LPS-treated microglia compared to unstimulated controls reveal shared and stimulus-specific responses, and stimulus-responsive markers. Color denotes condition- and direction-specific profiles as defined in the legend. (B) UpSet plot showing intersections among genes significantly upregulated or downregulated in Aβ and LPS treated microglia versus unstimulated controls. Intersection bars are colored according to the profile scheme shown in (A). (C) top row: GO term enrichment of shared, Aβ-, and LPS-enriched upregulated genes; bottom row: GO term enrichment of shared, Aβ-, and LPS-enriched downregulated genes. GO term plots are colored according to the profile scheme shown in (A). GO term labels were shortened for visualization; full pathway descriptions are provided in Table S4. Arrows indicate positive (↑) or negative (↓) regulation.

Commonly upregulated genes across both stimuli were enriched for inflammatory and chemotactic responses, including chemotaxis, CXCR chemokine receptor binding, cytokine-mediated signaling, colony-stimulating factor (CSF) signaling, lipid storage, and regulation of apoptotic pathways (Fig. 4C, top left panel). Aβ-stimulated iMGLs uniquely upregulated genes associated with Golgi lumen acidification, MAPK signaling, and apoptosis (Fig. 4C, top middle panel), while LPS-stimulated iMGLs exhibited strong activation of cytokine signaling, metal ion stress, protein refolding, and macrophage apoptotic processes (Fig. 4C, top right panel). These distinct transcriptional signatures indicate that Aβ and LPS drive partially overlapping but separable inflammatory and stress response programs in microglia.

Prompted by these findings, we further examined programmed cell death–associated pathways (Fig. S3A). Among the four major categories analyzed, apoptosis emerged as the most strongly induced in both Aβ- and LPS-treated conditions, with especially prominent induction in response to LPS. Genes associated with pyroptosis also showed moderate upregulation following LPS treatment, whereas necroptosis- and ferroptosis-related gene sets were more modestly induced. These results align with the transcriptional evidence for enhanced stress and cytotoxic signaling in response to these disease-relevant stimuli.

In parallel, downregulated DEGs shared across both treatments were significantly enriched in G protein–coupled signaling and T cell receptor activity (Fig. 4C, bottom left panel). Notably, LPS exposure led to a strong suppression of immune surveillance programs, including MHC class II receptor activity, antigen processing, and a microglial activation module (Fig. 4C, bottom right panel). However, the latter module consisted of only three genes—*TYROBP*, *GRN*, and *TREM2*—suggesting that this annotation may reflect a limited and selective component of microglial homeostasis rather than a broad suppression of activation. This more restrained signal may point to context-dependent regulation of immune homeostatic genes in LPS-activated microglia, rather than a global inhibition of immune function. Together, these findings offer a molecular framework for understanding how distinct pathological stimuli remodel microglial gene networks through both shared and unique immune regulatory signatures.

### Functional assays validate stimulus-specific stress responses in dCas9-KRAB iMGLs

To evaluate the functional outcomes of Aβ and LPS stimulation, we assessed multiple stress and immune markers in dCas9-KRAB iMGLs. Both Aβ and LPS induced a significant increase in lipid accumulation, as measured by LipidTOX staining (Fig. 5A), consistent with transcriptomic enrichment of lipid storage pathways. Aβ exposure also led to a significant elevation in mitochondrial oxidative stress, indicated by increased MitoSOX intensity, while LPS treatment showed a non-significant upward trend (Fig. 5A). In contrast, caspase-3 staining revealed only minimal, non-significant increases in both conditions (Fig. 5A), suggesting limited apoptotic cell death, in line with the absence of strong *CASP3* transcriptional induction (sFig. 3A).

**Figure 5.**
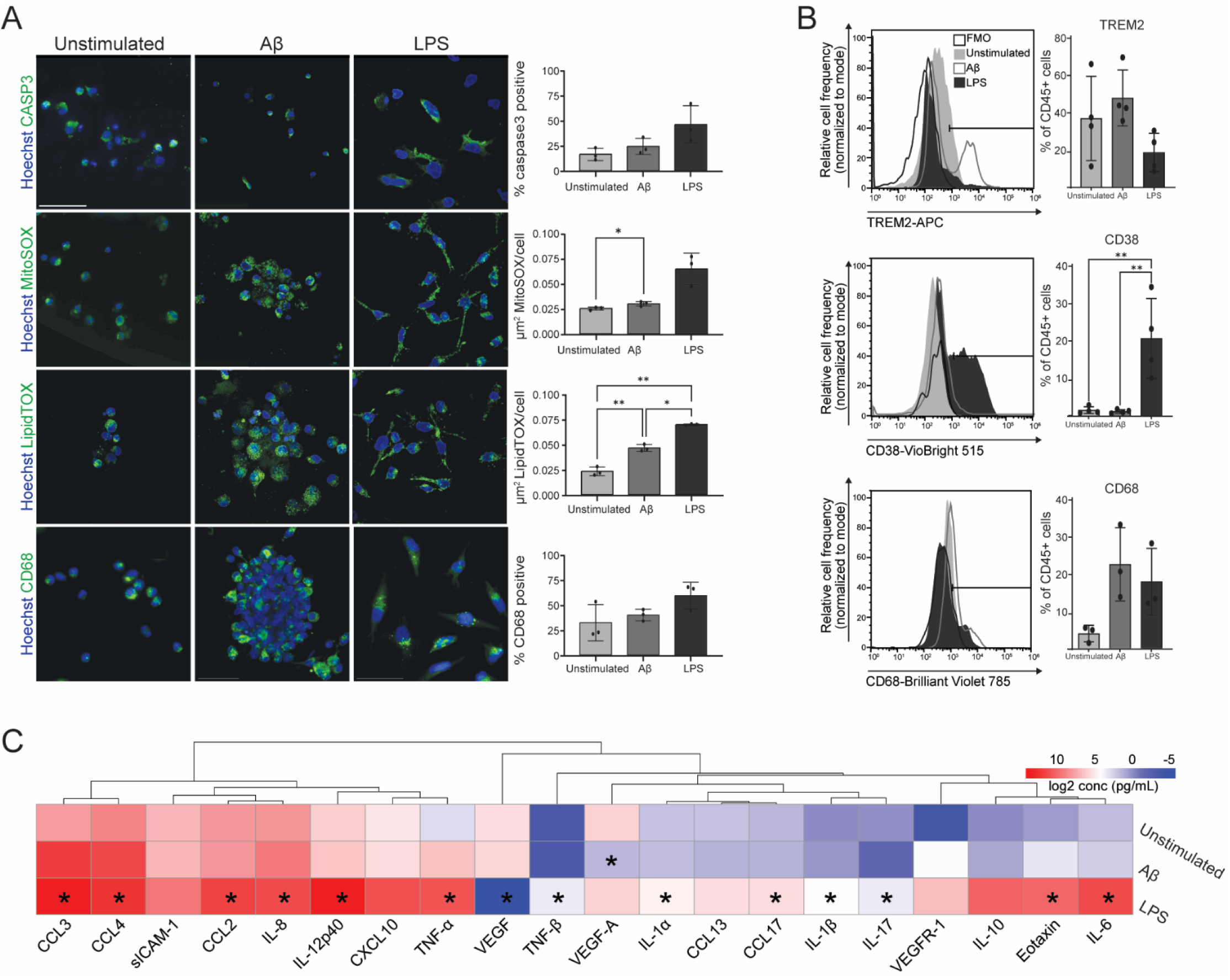
Cellular assays confirmed Aβ- and LPS-induced cellular stress and inflammatory responses. (A) Representative ICC images (scale bar: 50 µm) and quantification of caspase-3, MitoSOX, LipidTOX, and CD68 across treatment conditions. Data represent 3 independent iMGL differentiations, with ∼700–1,800 cells quantified per condition (mean ± SD). *\*p* < 0.05; \*\**p* < 0.01 (repeated-measures one-way ANOVA with Tukey’s multiple-comparisons test). (B) Flow cytometry analysis of TREM2, CD68, and CD38 expression across three conditions. Data represent mean ± SD. *\*p* < 0.05; \*\**p* < 0.01 (repeated-measures one-way ANOVA with Tukey’s multiple-comparisons test). (C) Cytokine profiling reveals Aβ- and LPS-induced cytokine/chemokine responses. Data represent 4 independent iMGL differentiations (mean ± SD) *\*p* < 0.05 (Student’s t-test comparing Aβ or LPS to unstimulated).

Flow cytometry showed significant upregulation of CD38 in LPS-treated cells, whereas TREM2 and CD68 levels remained largely unchanged, highlighting selective immune activation under LPS (Fig. 5B). Finally, multiplex cytokine profiling revealed distinct inflammatory signatures, with LPS driving robust secretion of pro-inflammatory mediators including CCL2/3/4, IL-6/8, and TNF-α (Table S5). Aβ elicited only mild, non-significant cytokine changes. Notably, VEGF and VEGF-A secretion were significantly reduced in the LPS and Aβ conditions, respectively, pointing to a suppression of trophic signaling (Fig. 5C).

Together, these results confirm that Aβ and LPS elicit partially overlapping but distinct stress programs in iMGLs, consistent with transcriptome-defined signatures of inflammation, metabolic stress, and limited apoptosis.

### Anchoring dCas9-KRAB iMGL cell states to in vivo microglia states and gene module signatures

To contextualize our in vitro model with known in vivo microglial diversity, we further subclustered the four major iMGL clusters into 13 transcriptionally distinct subclusters (Fig. 6A; Table S6). Each subcluster exhibited a unique gene expression profile, reflecting stimulus- and state-specific microglial programs (Fig. 6B).

**Figure 6.**
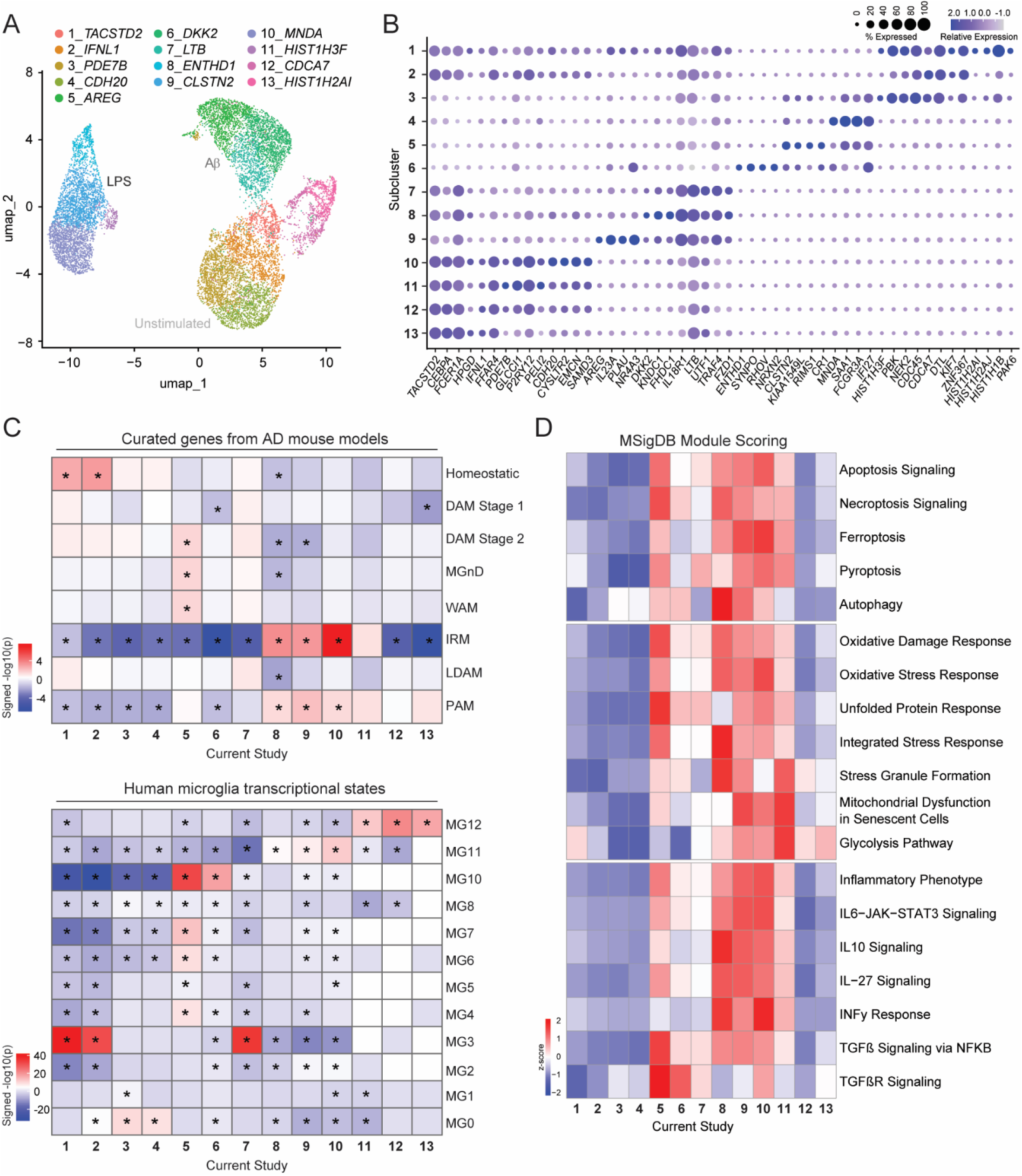
Subclustering reveals stimulus-specific microglial heterogeneity and links Aβ- and LPS-responsive states to in vivo microglia signatures. (A) UMAP visualization of subclustering results, showing 13 transcriptionally distinct microglial subclusters derived from the four major clusters identified in Figure 1. (B) Dot plot showing the top 4 differentially expressed genes for each subcluster. (C) Heatmap showing enrichment of curated mouse AD-associated genes and reported human AD-associated microglial states across subclusters. GSEA was performed and plotted values represent sign (NES) x −log10(adjusted *p*) for each signature. *\*p* < 0.05. (D) Heatmap showing module scores for selected MSigDB gene sets across microglial subclusters. Curated mouse genes for each model are provided in Table S6.

We first compared these subclusters to reference microglial states defined in mouse models (Clayton et al., 2021; Keren-Shaul et al., 2017; Li et al., 2019; Marschallinger et al., 2020; Roy et al., 2022; Safaiyan et al., 2021) (Fig. 6C). Unstimulated subclusters (subclusters 1-4), especially subclusters 1–2, were enriched for homeostatic microglia. In contrast, Aβ-associated subclusters (subclusters 5-7), particularly subcluster 5, showed strong enrichment for disease-associated microglia (DAM) stage 2, neurodegeneration-associated microglia (MGnD), and white matter-associated microglia (WAM). LPS-associated subclusters (subclusters 8-11) mapped more closely to interferon-responsive microglia (IRM) or proliferative-region-associated microglia (PAM), indicating a distinct inflammatory activation program induced by LPS exposure.

We next mapped iMGL subclusters to published human single-nucleus RNA-seq datasets from control and AD postmortem brains (Sun et al., 2023) (Fig. 6C). Unstimulated subclusters mapped to ribosome biogenesis or homeostatic microglia (MG3/MG0). Aβ-treated subclusters 5 and 7 strongly showed strong correspondence with inflammatory (MG10) and ribosomal (MG3) programs. In contrast, LPS-treated subclusters (8–11) displayed only modest enrichment for human microglial states, mapping weakly to antiviral (MG11) or cycling (MG12) profiles. Notably, proliferative subclusters 12 and 13, which emerged from the major proliferative cluster, aligned most strongly with cycling human microglia (MG12). Together, these results underscore both stimulus- and subcluster-specific differences in mapping fidelity, with Aβ-induced subclusters exhibiting closer resemblance to AD-associated human signatures than LPS-exposed subclusters.

To assess the functional consequences of Aβ and LPS stimulation, we performed module score analysis using curated gene sets from the Molecular Signatures Database (MSigDB) (Castanza et al., 2023; Liberzon et al., 2011, 2015; Subramanian et al., 2005) related to inflammation, stress response, and cell death (Fig. 6D; Table S6). Both Aβ and LPS stimulation induced increased cell death, stress response, and inflammatory gene expression across multiple subclusters. Subcluster 5 and subclusters 8–11 showed the greatest enrichment for oxidative stress, unfolded protein response, inflammatory signaling, and programmed cell death pathways. These coordinated transcriptional programs suggest a tight coupling between inflammatory activation and cytotoxic stress under pathological stimulation.

## Discussion

In this study, we establish that a commercially available human dCas9-KRAB iPSC line can be differentiated into iPSC-derived microglia-like cells (iMGLs) that recapitulate expected differentiation trajectories while retaining functional CRISPRi machinery (Fig. 1–2). We also establish that these iMGLs can mount distinct and measurable transcriptional and functional responses to acute inflammatory stimuli—amyloid-β (Aβ) and lipopolysaccharide (LPS)—commonly used to model Alzheimer’s disease–associated and classical inflammatory insults, respectively. Using single-cell RNA-seq integrated with morphological, functional, and cytokine-based assays, we demonstrate that both treatments induce overlapping inflammatory and chemotactic responses, including chemotaxis, CXCR chemokine receptor binding, cytokine-mediated signaling, colony-stimulating factor (CSF) signaling, lipid storage, and regulation of apoptotic pathways (Fig. 4) These shared features reflect conserved elements of the microglial response to environmental stress.

Despite these overlaps, Aβ and LPS triggered distinct transcriptional and cellular states. Aβ exposure led to a more selective and targeted gene expression program, prominently involving Golgi lumen acidification and MAPK signaling. In contrast, LPS exposure induced broader transcriptional remodeling across cytokine signaling, metal ion response, and protein refolding pathways (Fig. 4). Functionally, LPS-treated iMGLs showed stronger activation, with higher CD38 expression, elevated cytokine release, and increased lipid accumulation compared to Aβ-treated cells (Fig. 5). These findings highlight the distinct magnitude and breadth of inflammatory reprogramming induced by the two stimuli.

Even in the absence of stimulation, iMGLs exhibited a primed state enriched for immune activation (Fig. 3), consistent with previous observations that iPSC-derived microglia do not fully revert to a homeostatic baseline *in vitro,* and instead retain features of developmental or immune-ready states. (Abud et al., 2017; Butovsky and Weiner, 2018; Muffat et al., 2016). In our system, Aβ and LPS exposure further reshaped these baseline states, supporting the notion that iMGLs capture a flexible spectrum of microglial responses to environmental perturbation. To benchmark our system, we compared stimulus-exposed iMGL states to published in vivo microglial datasets. While not identical, both Aβ- and LPS-treated states showed partial convergence with disease-relevant activation signatures, underscoring the value of this *in vitro* model for capturing key features of microglial phenotypes (Fig. 6).

These results support the use of the dCas9-KRAB iMGL platform as a scalable, tractable system for modeling stimulus-specific microglial responses. Its compatibility with CRISPRi-based functional interrogation provides a foundation for future studies of gene–environment interactions and screening applications in neuroinflammation and neurodegeneration. Notably, the system leverages a publicly available AICS-0090 iPSC line and an established, widely adopted microglial differentiation protocol, enhancing its reproducibility and accessibility for the broader research community.

## Methods

### Maintenance of iPSCs

All stem cell work was performed in compliance with the Rush University Medical Center IBC committee. The dCas9-KRAB iPSC line was developed at the Allen Institute for Cell Science and available through the Coriell Institute for Medical Research (AICS-0090). This line was derived from the WTC parental line released by the Conklin Laboratory at the J. David Gladstone Institute. The AICS-0090 dCas9-KRAB iPSC line has been validated for genomic integrity, pluripotency, and differentiation into multiple lineages, as documented in the publicly available [Certificate of Analysis (CoA)](https://www.coriell.org/0/PDF/Allen/ipsc/AICS-0090-391_CofA.pdf) (Supplemental File 1). Upon receipt, the cells were expanded and cryopreserved into a working cell bank. For experiments, cells were thawed and cultured on Matrigel hESC-qualified matrix (Fisher Scientific, Cat# 08-774-552) in mTeSR™ Plus medium (StemCell Technologies, Cat# 05825). Cells were passaged every 3–4 days using ReLeSR (StemCell Technologies, Cat# 05872) and maintained at a split ratio of 1:3 prior to differentiation. Cultures were periodically monitored by karyotyping and tested monthly for mycoplasma contamination to ensure genetic and microbial stability. While we did not generate the cell line or establish the original master cell bank, we created a working cell bank of ∼20 vials and used cells that had undergone 3–4 passages prior to initiating the differentiation protocol.

### Generation of iPSC-derived microglia like cells (iMGLs)

iMGLs were generated according to previously published protocols (Abud et al., 2017; McQuade et al., 2018) with minor adjustments. iPSCs were seeded in Matrigel-coated 12-well plates (Fisher; 07-200-82) in mTESR Plus (STEMCELL Technologies; 100-0276) supplemented with Y-27632 ROCK Inhibitor (RI,10mM; STEMCELL Technologies, 72304) and cultured for 24-48 hours. On d0, colonies were switched to iPSC-derived hematopoietic progenitor (iHPC) basal media supplemented with FGF2, BMP4, Activin-A, and LiCl. On d2, cultures were fed with iHPC basal media supplemented with FGF2 and VEGF. On d4, cultures were fed with iHPC basal media supplemented with FGF2, VEGF, TPO, SCF, IL-6, and IL-3. Cultures were given a half media change every two days until d12. On d12, iHPCs were collected, washed, resuspended in complete iMGL differentiation media, and replated in Matrigel-coated 6-well plates (Fisher; 07-200-83). Cultures were supplemented with 1 mL fresh complete microglial differentiation media every other day and expanded every 4 days until d24. On d30, cells were expanded 1:2 in a 1:1 mix of complete differentiation media and complete maturation media. From d30–d40, cultures were supplemented every other day with 1 mL additional complete iMGL maturation media. Media formulations and concentrations are listed in Table S1.

### LPS stimulation

iMGLs were collected between d38 – d39 of the differentiation and counted. iMGLs were then re-plated in a minimum of 2 mL of conditioned maturation media on hESC-qualified Matrigel coated 6-well plates and allowed to incubate overnight. iMGLs were treated with LPS (1 ug/mL; Thermo, 00-4976-93) for 24 h. At the end of the 24 h incubation, plates were incubated on ice for 5 – 10 min to promote cell detachment of adherent cells, then collected and used for downstream applications.

### Aβ preparation and labeling

Amyloid-beta (Aβ1-42) was reconstituted and aggregated according to Labeling Amyloid Beta with pHrodo Red protocol from Fujifilm Cellular Dynamics Inc. All reagents required to reconstitute Aβ were provided in the aggregation kit (rPeptide, A-1170-2). This preparation yields aggregated Aβ species generated using a standardized labeling and aggregation protocol, without further biochemical characterization of aggregate composition.

### Aβ phagocytosis assay

iMGLs were collected between d40 – d42 of the differentiation and counted. iMGLs were then re-plated in a minimum of 2 mL of conditioned maturation media on hESC-qualified Matrigel coated 6-well plates and allowed to incubate overnight. pHrodo-labelled Aβ was thawed to room temperature and resuspended by vortexing. For all Aβ stimulation experiments, pHrodo-labeled Aβ aggregates were added to cultures at a final concentration of 25 µg/mL (1:200 dilution) and incubated for 2 h. Plates were then incubated on ice for 5 – 10 min to promote cell detachment of adherent cells, then collected and used for downstream applications.

### Flow cytometry

After stimulation, collected cells were used for flow cytometry. The buffer for Fluorescence-activated cell sorting (FACS) buffer was prepared by diluting 30% BSA (Fisher Scientific, 50-203-6474) in DPBS to a final 2% solution. Cells were pelted by centrifugation at 300 x g for 5 min at 4 °C, washed twice, and resuspended in the FACS buffer. Cells were then simultaneously stained with Human TruStain FcX according to the manufacturer’s instructions (Biologend; 422302) and with an appropriate fluorescent fixable live/dead dye (Thermo) on ice for 15 min. Fluorophore-conjugated antibodies or their isotype controls were incubated on ice for 30 min, then washed twice with FACS buffer. Stained cells were fixed with Fixation Buffer (Biolegend, 420801) according to the manufacturer’s instructions. If applicable, fixed cells were permeabilized using Intracellular Staining Permeabilization Wash Buffer (Biolegend, 421002) according to the manufacturer’s instructions and cells were stained for intracellular markers according to the above instructions. Data was collected using the Sony SH800 Cell Sorter and flow cytometry data was analyzed using FlowJo v10. Compensation metrics were set using unstained and single-color compensation controls and gating was determined using Fluorescence Minus One (FMOs) control. All live/dead dyes, conjugated antibodies, isotypes and their dilutions are listed in Table S1.

### Immunostaining and image acquisition

iMGLs were collected between d40–d42 of differentiation and replated overnight on Matrigel-coated Ibidi µ-Slide 8-Well high chamber slides (Ibidi, 80806) at a density of 50,000–75,000 cells per chamber. Cells were stimulated with LPS for 24 h or Aβ for 2 h and fixed with 4% paraformaldehyde (PFA; Electron Microscopy Sciences, 15714-S) for 15 min at room temperature. After three washes with DPBS, cells were permeabilized and blocked in 5% normal donkey serum (Jackson ImmunoResearch, 017-000-121) containing 0.02% Triton X-100 (Fisher, BP151100) for 1 h at room temperature.

Cells were incubated with primary antibodies diluted in blocking buffer for 2 h at room temperature, washed three times with DPBS, and incubated with secondary antibodies diluted in blocking buffer for 45 min at room temperature in the dark. For staining panels requiring two primary antibodies raised in the same species, the second primary antibody was conjugated to a 647-nm fluorophore using the Flexible CoraLite Plus 647 Antibody Labeling Kit (Proteintech; Rabbit IgG: KFA003, Mouse IgG1: KFA023, Mouse IgG2a: KFA043) according to the manufacturer’s instructions. Conjugated primary antibodies were incubated for 2 h at room temperature. Following antibody staining, cells were washed three times with DPBS and counterstained with Hoechst 33342 (ThermoFisher; 62249) diluted 1:2000 in DPBS for 3 min at room temperature. Cells were washed once more with DPBS and imaged. Primary and secondary antibodies and their dilutions are listed in Table S1.

### Morphological quantification

Live-cell phase-contrast images were acquired using a Nikon Ts2 inverted microscope and fixed immunofluorescence images were acquired using a Nikon Ti2-E widefield epifluorescence microscope, both operated using Nikon NIS-Elements imaging software. Morphological activation was quantified by measuring the proportion of cells bearing processes longer than 3 µm. Measurements were performed on both live-cell phase-contrast images and fixed immunofluorescence images using ImageJ. Cellular processes were defined as thin extensions emerging from the soma and exceeding the specified length threshold. For immunostained samples, only IBA1⁺ or CD45⁺ cells were included in the analysis. Data were derived from 3 independent iMGL differentiations. For each condition, 3-4 randomly selected, non-overlapping fields were analyzed per well at 10x magnification. The total number of cells quantified for each experiment is reported in the corresponding figure legends.

### Intracellular stress assay & quantification

LPS or Aβ stimulated iMGLs were stained for MitoSOX Red (Thermo, M36008) or LipidTOX Red (Thermo, H34476) was performed according to the manufacturer’s instructions. Cells were fixed with 4% PFA for 15 min at room temperature, washed three times with DPBS, and counterstained with Hoechst 33342 diluted 1:2000 in DPBS for 3 min at room temperature.

Cells were imaged using a Nikon AX/AX R with NSPARC confocal microscope and Nikon NIS-Elements imaging software. Quantification was performed by measuring positively stained area per field and normalizing to nuclei count to account for stimulus-induced changes. Data were derived from 3–4 independent iMGL differentiations. For each condition, 3-4 randomly selected, non-overlapping fields were analyzed per well at 10x magnification. The total number of cells included in the analysis is reported in the corresponding figure legends.

### Mesoscale multiplex biomarker assay

iMGLs were replated in 2 mL of conditioned maturation media overnight prior to stimulation with LPS or Aβ as described above. Media was collected and analyzed using the V-plex Human Biomarker 39-Plex Kit (Mesoscale Discovery; K15209D-1) according to the manufacturer’s protocol for each treatment condition from 4 independent iMGL differentiations.

### Prioritization of Alzheimer’s Disease-associated genes expressed in microglia

The prioritization of candidate genes was conducted using multiple criteria. These included the selection of representative genes from each AD GWAS loci, as reported in the EADB GWAS, along with genes with colocalization posterior probabilities (PP4) higher than 0.7, calculated using coloc, against multiple eQTL/sQTL, including bulk RNA-seq across various brain tissues (MSBB, ROSMAP, and Mayo Clinic), LCL RNA-seq, and sorted microglia RNA-seq (Bellenguez and EADB, 2020; Giambartolomei et al., 2014). Genes reported in another AD GWAS, with prioritization model probability higher than 0.5, and coloc results higher than 0.7 (across multiple eQTL datasets)(Schwartzentruber et al., 2021). Genes identified in the colocalization results reported in the MiGA study (Lopes et al., 2022), which included four AD GWAS datasets, along with transcriptome data from in-vivo microglia eQTL and ex-vivo microglia eQLT/sQTL from 4 brain regions (plus meta-analysis). And finally, genes reported by the microglia-specific eQTL and caQTL study that applied multiple prioritization methods, including coloc, moloc, ABC model, and overlap with microglia ATAC-seq peaks. Selected genes are listed in Table S2.

### Single-guide RNA (sgRNA) design, cloning and lentiviral library generation

For each identified gene, three sgRNA targets were chosen between −50 and +300bp of the transcriptional start site. Targets were chosen from top hits identified by CRISPick (Broad Institute) and E-crisp (Boutros lab). In addition to 165 genes prioritized for analysis, sgRNAs targeting control genes, CSF1R, CSF2RB, CDK8, and TGFBR2, previously identified as regulators of survival, as well as non-targeting controls were included. The vector pCRISPRia-v2 was a gift from Jonathan Weissman (Addgene 84832) (Dräger et al., 2022). This was modified to express eGFP amplified from pXR001, a gift from Patrick Hsu (Addgene 109049), in place of BFP by restriction digestion at the EcoRI and CsiI sites and T4 ligation (NEB). Guide RNA sequences were appended with appropriate 5’ and 3’ sequences, and a pool of oligonucleotides was synthesized (Twist Bioscience). The vector was digested with BstXI and BlpI, and oligonucleotides were amplified by PCR and subsequently cloned by Gibson assembly as previously described to generate an sgRNA library. Plasmids were amplified and prepared by MaxiPrep for lentiviral generation as previously described. Lentivirus was prepared by transfecting 293T cells in T75 flasks with the plasmid pool as well as psPAX2 and pVSVg (Gifts from the Trono and Reya labs, Addgene 12260 and 14888, respectively) helper plasmids using FuGene HD (Fisher, NC2132803).

### Sequencing and analysis of sgRNA distribution

Genomic DNA was amplified with barcoded next-generation sequencing (NGS) primers, adapted from previously published protocols (Joung et al., 2017), to include unique indices on both forward and reverse primers as well as a 10nt variable region just upstream of the 5’ vector binding sequence. PCR reactions from each sample were pooled, and DNA was purified with dual size selection with SPRI beads and analyzed on Fragment Analyzer. Sequencing libraries were diluted and loaded onto Illumina iSeq 100. For CRISPR screening analysis, BCL files were converted to FASTQ format. Raw reads were aligned to the reference with seqtk. Counts were then converted to phenotype scores with MAGeCK-iNC (Tian et al., 2019) modified so that two of the top three sgRNA sequences per gene were analyzed. With this pipeline, comparisons between adjacent time points day 12, day 24, day 36 and day 45 were made to assess the effect of gene knockdown at different stages of iMGL differentiation and maturation (Table S2). Heatmap displaying genes with significant changes in abundance in at least one of these three intervals was made withggplot2 (v3.5.1) (Wickham, 2009) in R. Epsilon reflects abundance phenotype such that positive numbers reflect a reduction in corresponding gene-targeting cell enrichment.

### CRISPRi survival screen

dCas9-KRAB iPSCs were dissociated in Accutase (Thermo; 00-4555-56) and seeded into 6 well tissue culture-treated plates coated with Matrigel. Cells were transduced with lentivirus at low MOI to generate cultures with 30% positive eGFP signal and an average of at least 1000x sgRNA coverage. Cells were cultured overnight in the presence of 10µM Y-27632 and mTeSR Plus. Media was changed the following day and cells were selected starting the next day with 2 µg/ml puromycin (Fisher; AAJ67236XF), for a duration of four days and allowed to recover without selection. Cells were then seeded as small colonies in 15cm dishes for HPC differentiation and subsequent iMGLs differentiation as described above. Cells were collected and pelted at days 0, 12, 24, 36 and 45 of differentiation. Pellets were subjected to gDNA extraction (Zymo).

### Sample preparation, library construction, single-cell RNA sequencing, and data processing

Single-cell RNA sequencing experiments were performed using cells derived from a single iMGL differentiation, with unstimulated, Aβ-treated, and LPS-treated cells processed in parallel. Single-cell suspensions were processed into sequencing-ready libraries using the 10x Genomics Chromium Single Cell 3′ Platform. Libraries were sequenced on a NovaSeq 6000 (Illumina) at a depth of 50k reads/cell. Following sequencing and FASTQ generation, raw count matrices were produced using CellRanger v6.0.1. RNAs were mapped to the GRCh38 reference genome via STAR. Transcripts were assigned to individual cells and duplicate reads removed. The resulting count matrices were processed using the Seurat (v5.3.0) package (Hao et al., 2024) in R. We applied quality controls metrics to the single-cell RNA sequencing results to remove cells with poor transcriptomic quality (nCount_RNA >7500 & < 70,000; nFeatures_RNA >2500 & < 8,000; percent.mt <10%; percent.rb to <30%). Seurat::SCTransform was used to normalize and scale the UMI counts for each individual library based on regularized negative binomial regression. Principal component analysis was performed, followed by Seurat::FindNeighbors (dims = 1:10) and Seurat::FindClusters (res = 0.75) to identify clusters based on a shared nearest neighbor clustering algorithm. Data was visualized with Seurat::RunUMAP(dims = 1:15). The data was inspected for cell doublets using DoubletFinder (v2.0.4) (Hao et al., 2024; McGinnis et al., 2019) and those containing mixed cluster signatures were removed. Cell clusters were reprocessed using the functions described above prior to in-depth analysis.

### Marker gene identification and ClueGO cluster annotation

Differentially expressed genes (DEGs) were identified using Seurat::FindAllMarkers. For broad cluster comparisons (4 main clusters), a minimal effect-size filter was applied (logfc.threshold = 0.1, min.pct = 0.05, only.pos = FALSE) to capture cluster-defining transcriptional differences (Table S3). For higher resolution subcluster profiling, thresholds were adjusted (logfc.threshold = -inf, min.pct = 0.15, only.pos = FALSE) to enable detection of biologically relevant markers in smaller cell populations (Table S6). DEGs were filtered to remove non-coding transcripts prior to downstream analyses. Cluster and subcluster dot plots of the top DEGs for each transcriptional cluster were generated using ggplot2.

Gene Ontology (GO) network analysis for the four main clusters was constructed using ClueGO (v2.5.10) plugin in Cytoscape (v3.10.3) on the top 200 significant DEGs per cluster, both sorted by descending order of log2 fold change. ClueGO analysis was performed using the GO_BiologicalProcess-EBI-UniProt-GOA-ACAP-ARAP database (release 25.05.2022, 00:00). A two-sided enrichment/depletion test was used to determine ontology enrichment with Bonferroni stepdown correction (p ≤ 0.01, Table S3). Parameters for network constructions were as follows: GO levels 3 – 8 with GO term fusion to reduce redundant terms; Kappa score = 0.4; minimum genes included in term = 3; minimum percentage of genes included in term = 4%.

### Comparative transcriptional analysis and Gene Ontology (GO) enrichment analysis

To quantify stimulus-associated transcriptional differences, Seurat::FindMarkers was used to compare the Aβ-enriched (Cluster 1) and LPS-enriched (Cluster 2) transcriptional clusters to the unstimulated-enriched cluster (Cluster 0) (logfc.threshold = 0.1, min.pct = 0.05). DEGs were filtered to remove non-coding transcripts prior to downstream analyses. Filtered results were categorized into transcriptional response groups based on the direction and significance of their expression changes, including shared upregulated genes, shared downregulated genes, Aβ-enriched genes, LPS-enriched genes, and anti-correlate genes exhibiting opposing regulation between conditions (Table S4). Pearson correlation analysis was performed to quantify the relationship between Aβ- and LPS-associated log_2_ fold change values across all tested genes.

For functional pathway analysis, gene sets corresponding to shared and condition-specific transcriptional categories were analyzed separately, with all expressed genes in the dataset used as the background universe. Gene Ontology (GO) enrichment analysis was performed usingclusterProfiler::enrichGO (v4.14.3) (Yu et al., 2012), testing for enrichment across Biological Process, Molecular Function, and Cellular Component ontologies (ont = “ALL). Enrichment analyses were conducted using a *p* value cutoff of 0.05, minimum gene set sizes of 1–10 (depending on gene category), and a maximum gene set size of 500 (Table S4). Results were visualized using ggplot2.

### Comparison to mouse and human microglial gene state sets and module scoring

To compare iMGL subclusters to published mouse and human microglial state signatures (Clayton et al., 2021; Keren-Shaul et al., 2017; Li et al., 2019; Marschallinger et al., 2020; Roy et al., 2022; Safaiyan et al., 2021; Sun et al., 2023), differential expression statistics were used as continuous input for gene set enrichment analysis (GSEA). Genes were ranked using a score calculated as log_2_ fold change weighted by −log₁₀(adjusted p value). GSEA was performed separately for each iMGL subcluster using the mouse or human microglial gene sets, and enrichment results were considered significant at adjusted p < 0.05. For visualization, enrichment results were summarized by plotting sign (NES) × −log₁₀(adjusted p value) for each subcluster–gene set comparison and heatmaps were generated using ggplot2. Module scoring was performed using selected gene sets from the Molecular Signature Database (MSigDB) related to cell death, cellular stress, and inflammatory signaling (Table S6). Per-cell scores were calculated using Seurat:AddModuleScore and results were visualized using heatmap.

## List of abbreviations

iPSC: induced pluripotent stem cell
CRISPRi: CRISPR interference
Aβ: amyloid beta
LPS: lipopolysaccharide
AD: Alzheimer’s Disease
IHPCs: human induced pluripotent stem cell-derived hematopoietic progenitors
iMGLs: human induced pluripotent stem cell-derived microglia-like cells
GO: Gene Ontology
DGE: differential gene expression
DEGs: differentially expressed genes
DAM: disease-associated microglia
MGnD: neurodegeneration-associated microglia
WAM: white matter-associated microglia
IRM: interferon-responsive microglia
LDAM: lipid droplet-associated microglia
PAM: proliferative-region-associated microglia
MSigDB: Molecular Signatures Database
GPCR: G-protein-coupled receptor
PCD: programmed cell death
FACS: fluorescence-activated cell sorting

## Declarations

None

## Availability of data and materials

We generated raw and normalized count matrices of the single-cell RNA sequencing and CRISPR interference screen datasets from the dCas9-KRAB line. These data are available at GEO under accession code GSE306200 & GSE306202.

## Competing interest

The authors declare that they have no competing interests

## Funding

This study was supported by NIA grants R01AG074082 and R01AG079223 (to Y.W.), P30AG10161, P30AG72975, R01AG015819, R01AG017917, U01AG61356 and the Paul M Angell Family Foundation (to D.A.B.).

## Author contributions

E.M.K. and D.M.S. established the iMGL differentiation protocol in-house based on published methods. D.M.S. performed cellular assay validation experiments and conducted single-cell RNA sequencing data analysis. B.N. contributed to microglial state mapping analysis. N.A.K. and H.V. assisted with assay development. F.S. and R.A.V. contributed to early-stage functional screening efforts that supported the development of the study. S.D.T. conducted single-cell RNA sequencing experiments, and J.X. performed data processing. D.M.S. and Y.W. drafted, revised, and finalized the manuscript. D.A.B. provided insightful and critical feedback. Y.W. conceived and directed the study, led the research activities, and supervised manuscript preparation. All authors reviewed and approved the final manuscript.

## Acknowledgments

We acknowledge the support of all RADC staff for their assistance throughout this project. This research was made possible through the Allen Cell Collection, available from the Coriell Institute for Medical Research.

## Authors’ Information

Further information and requests may be directed to the corresponding author Dr. Yanling Wang

## Materials Availability

This study did not generate new unique reagents.

## Supplementary Figures

**Supplementary Figure 1.**
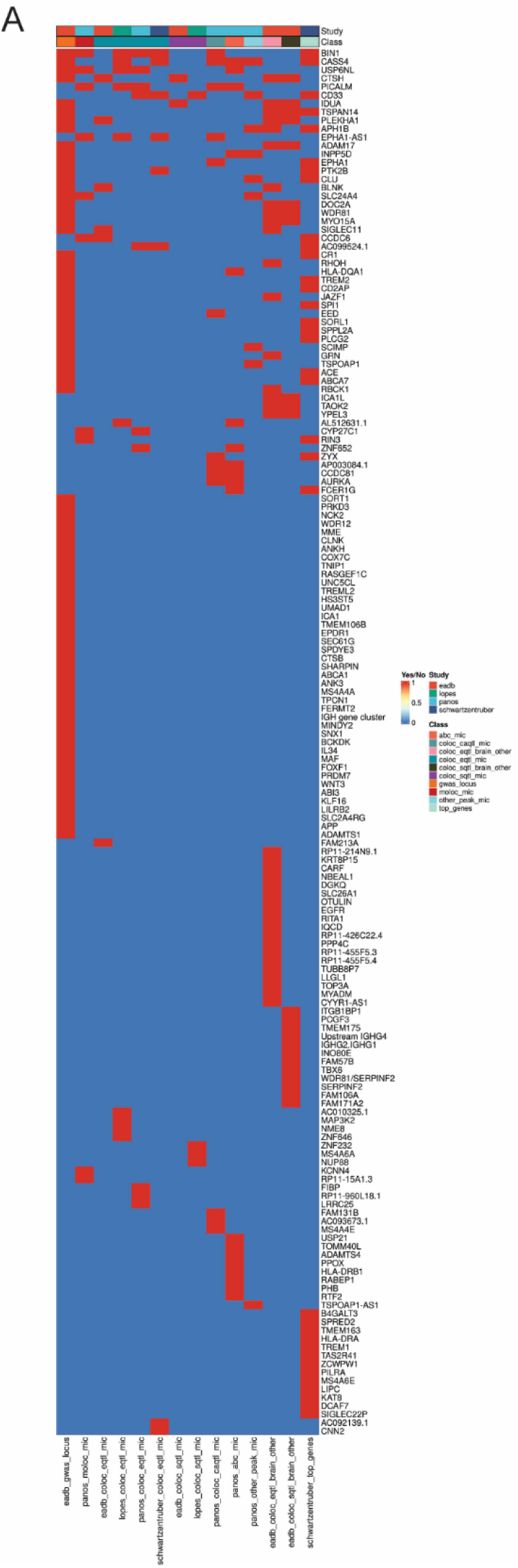
Origin and classification of CRISPRi pooled screen genes. (A) Heatmap showing the source study and class of the 165 genes included in the CRISPRi pooled screen.

**Supplementary Figure 2.**
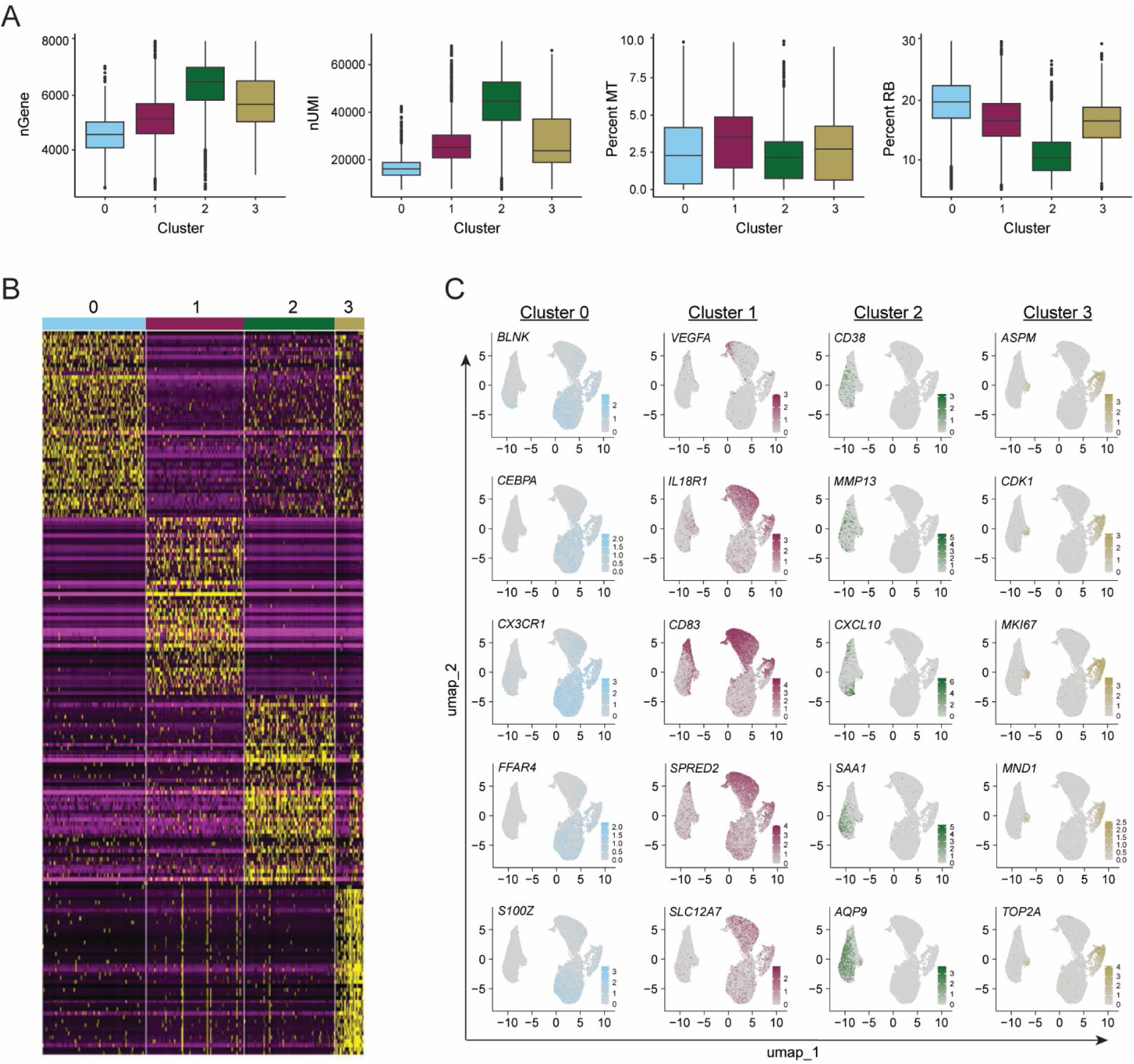
Single-cell RNA-seq quality control and microglial marker gene expression. (A) Boxplots showing quality control metrics across individual clusters, including number of detected genes (nGene), number of UMIs (nUMI), percentage of mitochondrial reads (percent.mt), and percentage of ribosomal reads (percent.RB). (B) Heatmap displaying the top 50 differentially expressed genes defining each cluster. (C) Feature plots illustrating the expression of microglial marker genes across clusters.

**Supplementary Figure 3.**
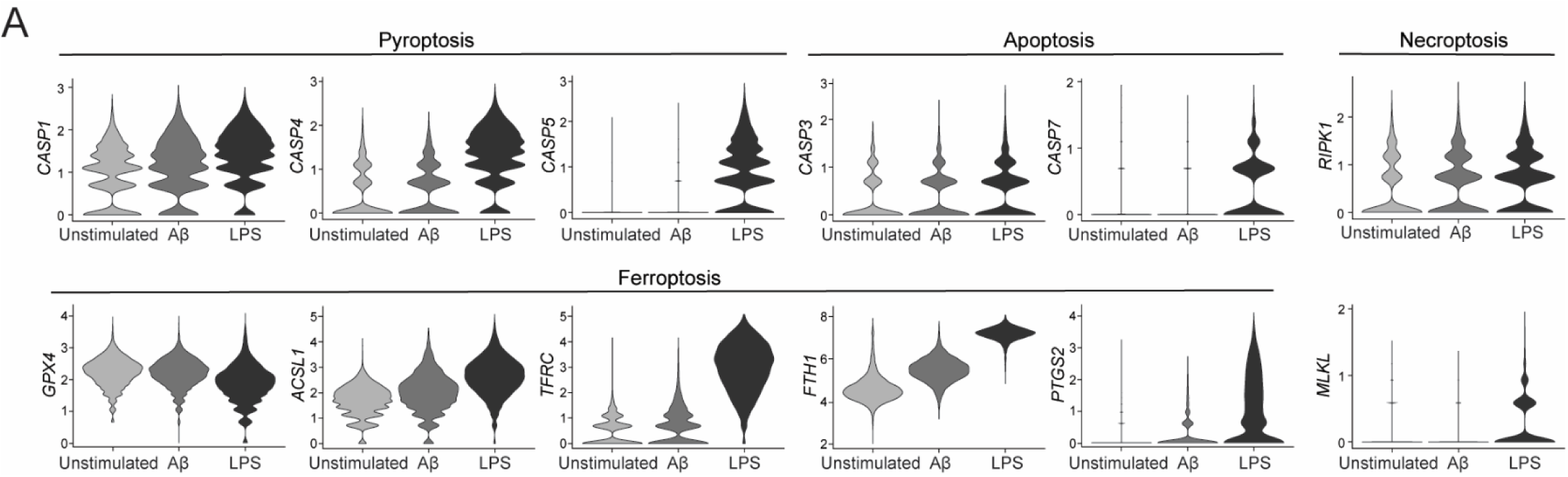
(A) Violin plots showing expression of genes related to pyroptosis *(CASP1*, *CASP4*, and *CASP5*), apoptosis (*CASP3* and *CASP7*), ferroptosis (*GPX4, ACSL4*, *TFRC*, *FTH1*, and *PTGS2*) and necroptosis (*RIPK1, MLKL*), each of which exhibited increased or decreased expression in Aβ- or/and LPS-treated microglia relative to unstimulated controls.

## Supplementary Tables

**Table S1.** Media and Assay Reagents

**Table S2.** Candidate Survival Genes from CRISPRi Screens

**Table S3.** iMGL Markers by Cluster and ClueGO Pathways

**Table S4.** Stimulus Specific DEGs and GO Terms

**Table S5.** MesoScale Cytokine Concentrations

**Table S6.** iMGL Markers by Subcluster and MSigDB Gene Sets

## Notes

### Competing Interest Statement

The authors have declared no competing interest.

### Summary of Updates

This revised version reflects significant updates from the original manuscript. Specifically, we added new experimental data in Figure 1 to strengthen the characterization of microglial differentiation. We also reshaped the overall narrative to better emphasize stimulus-specific responses and functional benchmarking. Several data analyses were re-performed to improve clarity and robustness, and new figures were added to support key conclusions. These changes enhance the reproducibility, accessibility, and translational relevance of the model, in alignment with the manuscript's focus as a resource.

